# Targeted metagenomics using probe capture detect a larger diversity of nitrogen and methane cycling genes in complex microbial communities than traditional metagenomics

**DOI:** 10.1101/2022.11.04.515048

**Authors:** H.M.P. Siljanen, L. Manoharan, A.S. Hilts, A. Bagnoud, R.J.E. Alves, C.M. Jones, M. Kerou, F. L. Sousa, S. Hallin, C. Biasi, C. Schleper

**Affiliations:** Department of Environmental and Biological Sciences, University of Eastern Finland, Kuopio, Finland; Department of Functional and Evolutionary Ecology, University of Vienna, Vienna, Austria; Department of Forest Mycology and Plant Pathology, Sweden University of Agricultural Sciences, Uppsala, Sweden; Department of Ecology, University of Innsbruck, Innsbruck, 6020, Austria

**Keywords:** metagenomics, nitrogen cycling, methane cycling, probe hybridization targeted metagenomics, PCR amplicon sequencing, shotgun metagenomics

## Abstract

Microorganisms are key players in the global cycling of nitrogen (N) and carbon (C), controlling their availability and fluxes, including the emissions of the powerful greenhouse gases nitrous oxide (N_2_O) and methane (CH_4_). Characterizing the microbial functional guilds driving these processes is crucial for understanding ecosystem functioning and predicting their responses to environmental changes. Standard sequence-based characterization methods often reveal only a limited fraction of their diversity in nature because of their low relative abundance, the insufficient sequencing depth of traditional metagenomes of complex communities, and limitations in coverage of PCR-based assays. Here, we developed and tested a targeted metagenomics approach based on probe capture and hybridization to simultaneously characterize the diversity of multiple key metabolic genes involved in inorganic N and CH_4_ cycling. We designed comprehensive probe libraries for each of the 14 target marker genes comprising 264,000 unique probes. These probes were used to selectively enrich the target genes in shotgun metagenomic libraries. In validation experiments with the mock communities of known microorganisms, targeted metagenomics yielded gene profiles similar to those of the original communities. Only GC content had a small effect on probe efficiency, as low GC targets were less efficiently detected than those with high GC, within the mock communities. Furthermore, the relative abundances of the marker genes obtained using targeted or traditional shotgun metagenomics from agricultural and wetland soils were significantly correlated, indicating that the targeted approach did not introduce significant quantitative bias. In addition, using archaeal *amoA* genes as a case-study, targeted metagenomics identified substantially higher taxonomic diversity and a larger number of sequence reads per sample, yielding diversity estimates 28 or 1.24 times higher than shotgun metagenomics or amplicon sequencing, respectively. Notably, shotgun metagenomics detected only three out of the 84 *amoA* gene phylotypes detected using targeted metagenomics. Our results show that targeted metagenomics complements current approaches to characterize key microbial populations and functional guilds in biogeochemical cycles in different ecosystems, enabling more detailed, simultaneous characterization of multiple functional genes.

**Manuscript contribution to the field:** Metagenomic sequencing often yields limited numbers of sequences of rare microbial taxa or functional genes, preventing in-depth analyses of specific populations and functional groups. Amplicon-based approaches enable the higher diversity coverage of target populations, but the drawback is the difficulty in designing unbiased primers that cover the highest intra-group diversity. Targeted metagenomics overcomes these challenges and results in similar community structure as traditional amplicon sequencing, while expanding the sequence space in a less biased metagenomic-based approach. Therefore, targeted metagenomics is an invaluable tool for studying the diversity of specific populations within complex natural microbiomes. Here, we present and evaluate a probe library designed for targeted metagenomics of nitrogen and methane cycling genes in complex communities.

## Introduction

The global nitrogen (N) and carbon (C) cycles are essential processes of the Earth’s biosphere and crucial for ecosystem functioning (Zaehle 2013, Gruber and Galloway, 2008; Archer 2010; Falkowski et al., 2000; Steffen et al. 2015). All major N transformation processes (i.e., nitrogen fixation, nitrification, denitrification, dissimilatory nitrate reduction to ammonium (DNRA), and anaerobic ammonium oxidation, or anammox), are performed exclusively by the functional guilds of bacteria, archaea, and eukaryotes (Galloway et al., 2004; Stein and Klotz, 2016; Kuypers et al., 2018; Offre et al. 2013). Their activities regulate N availability for primary producers and microorganisms across ecosystems and control the production and consumption of the potent greenhouse gas nitrous oxide (N_2_O) and other gaseous N compounds through processes such as nitrification, denitrification, and non-denitrifier N_2_O reduction (Barnard, et al., 2005, Thompson et al., 2012; Jones et al., 2014; Prosser et al. 2020). Methane (CH_4_) is another major product of the microbial trophic chain underlying C cycling, and, like the most inorganic N compounds, its production and consumption are also regulated by highly specific groups of microorganisms (Bodelier and Laanbroek 2004, Murrel and Jetten 2009, Conrad 2009). Importantly, CH_4_ is the second most potent greenhouse gas after carbon dioxide (CO_2_) and, together with N_2_O, contributes at least 25% of the total global warming caused by greenhouse gases (Myhre et al., 2013; Friedlingstein et al., 2020; Lan et al., 2023).

Despite their important ecological role, microorganisms participating in inorganic N and CH_4_ transformations typically constitute a small fraction of microbial communities in most soil, sediment, and aquatic ecosystems (Håvelsrud, et al., 2011; Palmer et al., 2012; Aanderud et al., 2015, Nelson et al, 2016, Ouygang and Norton 2020). Community profiling based on 16S rRNA genes only allows for very limited information about microbial functions (Alteio et al. 2021). Even in the rare cases when functions can be inferred from organism phylogeny, the striking genetic and functional diversity within functional guilds remains concealed, such as that among their key metabolic enzymes (e.g., ammonia monooxygenase (*amoA*), Alves et al. 2018; methyl co-enzyme M reductase (*mcrA*), Borrel, 2019). Additionally, taxonomy-based approaches are inadequate in elucidating processes that have a broader taxonomic distribution than originally thought (e.g. Kuypers et al. 2018, Borrel 2019, Li et al. 2023; Saghaï et al., 2024). Shotgun metagenomics is the typical method of choice for obtaining a relatively unbiased picture of the natural microbiome, provided that issues with sample preparation, DNA extraction, library preparation, and sequencing method can be ruled out (Pan et al., 2010, Nnadozie, Lin and Govinden, 2015; Sinha et al., 2017; Sui et al., 2020). Nevertheless, in complex and diverse microbial communities such as those in soils and sediments, even deeply-sequenced metagenomes do not uncover the full diversity of microbial functional guilds and key functional genes owing to their typically low relative abundances within the broader community (Palmer et al., 2012, Orellana et al. 2019, Ouygang and Norton 2020, Zhao et al. 2023). As a low-cost alternative, the characterization of specific functional groups has long relied on the PCR amplification of genes that encode key metabolic enzymes, as for example, of organisms involved in various N and CH_4_ cycling pathways (Knief 2015; Brauer et al. 2021; Alves et al. 2018; Romdhane et al. 2021, Clark et al. 2022). However, the gene diversity captured by gene-specific PCR assays is limited by the primers used, which introduce biases that make comparisons among different genes or samples difficult (Hallin and Lindgren, 1999; Throbäck al. 2004; Kolb et al., 2003; Siljanen et al., 2011). Furthermore, in complex microbial communities, target genes of interest are encoded by a large diversity of organisms, and usually include a large fraction of novel gene sequence variants from uncharacterized organisms (Jones et al. 2014; Saghaï et al 2023). Thus the geometric expansion of genomic data has made it increasingly obvious that designing a single PCR primer pair to target the inherent diversity of metabolic genes is highly problematic (Bonilla-Rosso et al. 2016). Therefore, linking ecosystem functions to microbial communities is often only possible through the complex and resource-intensive combinations of meta-omics and isotope labelling approaches (Tveit et al. 2015, Orellana et al. 2019).

Targeted high-throughput sequencing approaches, known as probe capture, also called hybridization capture, hybridization-based target enrichment, or captured metagenomics, have been used to study complex eukaryotic samples such as human exons, ancient human genomes, plant transcriptomes, and cancer marker single nucleotide polymorphisms (Skoglund et al. 2014, Ichida et al. 2019, Futema et al. 2012, Gasc et al., 2016, Kamil et al. 2021, Bewicke-Copley et al. 2019). A few studies have also used this approach to facilitate the in-depth study of microbial communities using either 16S rRNA genes or other microbial functional genes (Denonfoux et al., 2013; Manoharan et al. 2015, Noyes et al., 2017, Beaudry et al., 2021). The probe capture approach relies on targeting specific short sequences within a broader genomic pool (e.g., full genomes, genome or gene fragments) with biotin-labelled probes. The probes that hybridize with their target regions are then selectively captured from the full genomic library using streptavidin-labelled magnetic beads (Liu et al., 2016). Different from sequence capture, where multiple probes are designed to cover a whole exon, captured metagenomics relies on the large databases of multiple target genes clustered at predefined similarity cut-offs, which are used to design a large number of generic probes that are able to capture an extended sequence space for each of these sequence clusters (Kushwaha et al., 2015, Manoharan et al., 2015).

To improve detection and the characterization of important, but low abundant microbial guilds involved in N and CH_4_ cycling in natural communities, we developed and evaluated a probe-based targeted metagenomics approach to key genes involved in these processes. In addition to the enhanced resolution and coverage of genetic sequence space, this approach allows the parallel analysis of distinct steps within N- and CH_4_-cycling pathways, thus combining the advantages of both shotgun metagenomics and amplicon-based approaches. Here, we present a new probe set for targeting 14 distinct functional genes involved in inorganic N cycling (i.e., N fixation, nitrification, denitrification, anammox, and DNRA), and three genes involved in CH_4_ production and consumption. We evaluated our approach using mock communities comprising of DNA from microorganisms involved in inorganic N cycling, or CH_4_ production or consumption with varying gene GC% content, which is known to affect probe hybridization efficiency. As a proof-of-concept for complex communities, we performed shotgun metagenomics and targeted metagenomics on agricultural and wetland soil samples and compared the diversity of two functional markers, the archaeal *amoA* genes (marker for ammonia oxidizers) and *nosZ* genes (marker for nitrous oxide reduction).

## Materials and methods

### Construction of the target gene databases (TDBs)

Target gene databases (TDBs) were constructed, containing all identifiable variants for the following key genes: the nitrogenase iron subunit (*nifH*), bacterial and archaeal ammonia monooxygenase subunit A (*amoA)*, nitrite oxidoreductase beta subunit (*nxrB)*, hydrazine oxidoreductase A (*hzoA)*, formate dependent nitrite reductase (*nrfA*), periplasmic nitrate reductase alpha subunit (*napA*), respiratory nitrate reductase alpha subunit (*narG*), copper-containing nitrite reductase (*nirK*), cytochrome cd_1_ nitrite reductase (*nirS*), nitric oxide reductase subunit B (*norB*), nitrous oxide reductase (*nosZ*), particulate methane monooxygenase subunit A (*pmoA*), soluble methane monooxygenase component A alpha chain (*mmoX*), and methyl-coenzyme M reductase I subunit alpha (*mcrA*).

Hidden Markov Model (HMM) models were generated in order to identify all variants of the target genes from public databases (Fig. 1). These models were built based on reference sequence alignments from curated databases already available for selected genes, such as *amoA* (Alves et al., 2018), *pmoA* (Knief et al., 2015), *nosZ* (Jones et al., 2013), *nirK, nirS, nor* (Graf et al., 2014); and the Fungene repository (Fish et al., 2013). For target genes where alignments were not available, reference alignments were generated from gene sequence data publicly available on the NCBI WGS-database, from the full length open-reading frame of each subunit, to cover all known diversity in each gene. Structure-based searches of the Genbank nt- and envnt-database were subsequently performed with nhmmer using the generated HMMs for every target gene (Wheeler & Eddy, 2013) on a local supercomputer cluster (Centre of scientific computing CSC, Espoo, Finland) (July 2017). The HMM models are available through Zenodo: https://doi.org/10.5281/zenodo.15752134 (In script: probe-capture/2-selected_outputs/hmmer_profiles.zip). This sequence search selection process generated ∼600,000 unique sequences across all gene families. The obtained database was clustered to 100% identity in order to remove duplicates sequences in database with cd-hit (Fu et al., 2012) and manually inspected to exclude 16S rRNA genes. The final output comprised the target gene database (TDB; available through Zenodo: https://doi.org/10.5281/zenodo.15752134;inscript:probe-capture/2-selected_outputs/all_genes_compiled_ncbi_nt_envnt_fungene_split_file_0.fasta.gz-file (four files ..file_0-3.fasta.gz). The workflow can be seen in Fig 1. Scripts for Target gene database search is available in Zenodo: https://doi.org/10.5281/zenodo.15752134;/probe-capture/tree/main/3-other_scripts/Henri_scripts/Script_probe_capture.sh-folder.

**Fig. 1.**
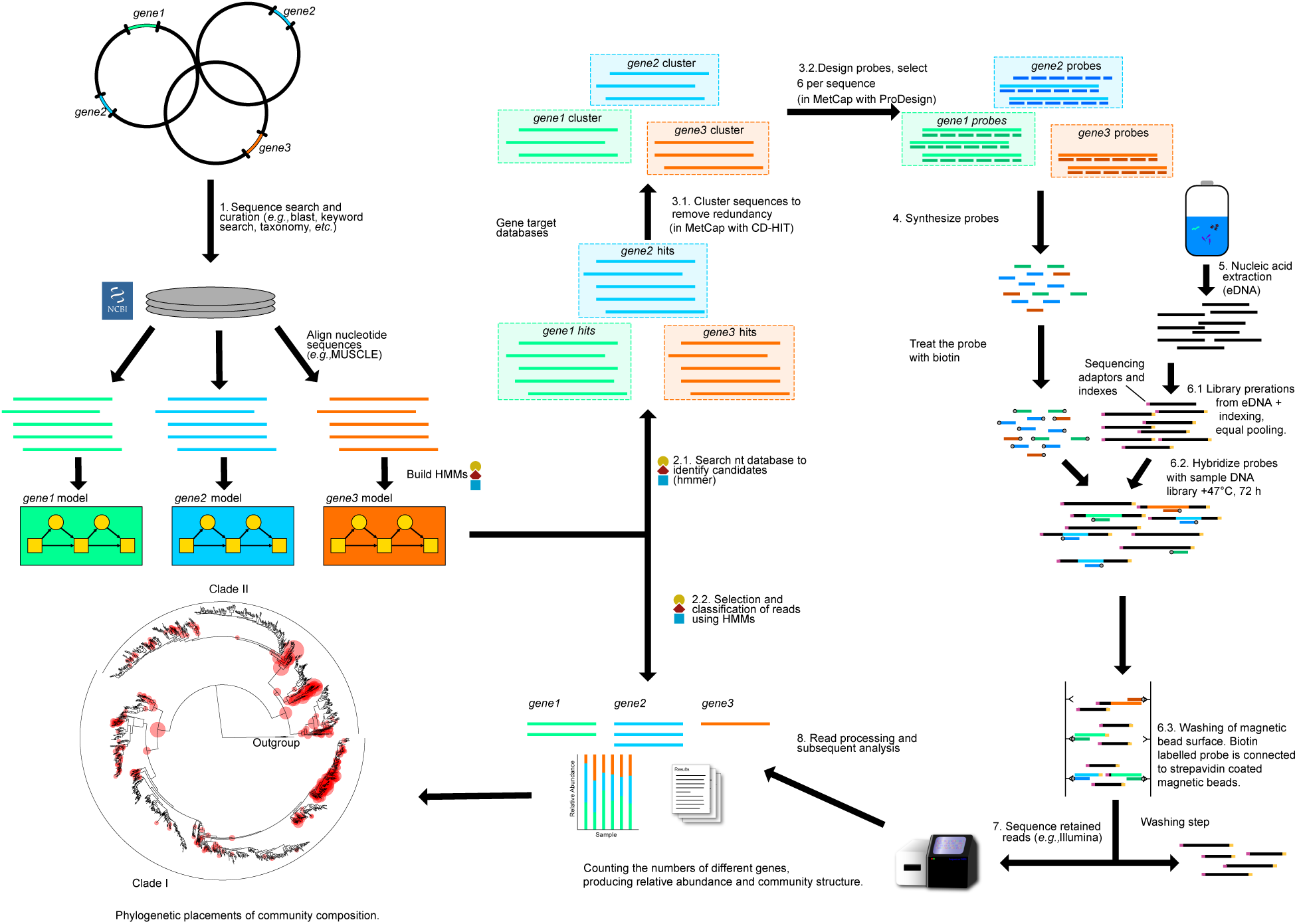
Illustration of the pipeline for the generation of HMM models for each gene, and how these models are used to study community composition with targeted metagenomics: 1. Steps followed for the generation of sequence databases for the genes of interest and subsequent building of the respective HMM models for each gene; 2.1. The produced HMM models were used to recruit more gene variants from the NCBI nt database and generate a target gene database for probe design; 2.2. The same HMM models were also used to classify the reads captured downstream; 3.1. Clustering of the target gene database with CD-HIT (in MetCap pipeline); 3.2. Design and selection of six 50-mer probes for each sequence cluster (in MetCap pipeline); 4. Synthesis and biotinylation of the designed probes by Roche; 5. DNA extraction of the sample of interest; 6.1. Sequencing libraries preparation and addition of sequencing adaptors and indexes; 6.2. Hybridization of the probes to indexed DNA libraries for 72 hours at +47°C; 6.3. Purification of the DNA hybridized to the probes with streptavidin coated magnetic beads; 7. Sequencing of the hybridized libraries with Illumina MiSeq; 8. Read processing with HMM profiles and taxonomy assignment for the genes of interest.

### Generation of probes for target genes

The MetCap bioinformatics pipeline was used as a protocol to produce probes from highly complex datasets for targeted metagenomics (Kushwaha et al., 2015). Default parameters were used for designing up to six unique 50mer probes for each sequence cluster in the TDBs (clustered with an 80% identity clustering threshold), with melting temperature of 47 °C, resulting in a set of 263,111 unique gene-specific probes (Fig 1, and in Zenodo: https://doi.org/10.5281/zenodo.15752134;probe-capture/2-selected_outputs/Final_N_probes.list1.fasta-file. The probes were synthesized by NimbleGen SeqCap EZ (Roche NimbleGen, Inc., Madison, USA) as Custom design (since the MetCap pipeline was used for probe design, probe order was done with Roche HyperDesign tool: https://www.hyperdesign.com), with biotin labelling to enable retrieval of the hybridized targets using streptavidin coated magnetic beads (Kushwaha et al., 2015; Manoharan et al., 2015).

### Extraction of DNA from cultures and generation of mock community DNA samples

DNA samples from mock communities encoding the genes of interest were generated by mixing genomic DNA from different organisms in variable proportions, in order to simulate variation in overall GC% content. The pool of DNA in the mock communities included DNA from: *Nitrosospira multiformis*, *Nitrososphaera viennensis*, *Nitrospira defluvii*, Ca. *Kuenenia stuttgartiensis* PCR fragment of the full-lenght *hzoA* gene, *Pseudomonas aeruginosa*, *Escherichia coli*, *Shigella sonnei*, *Cupriavidus metallidurans*, *Cupriavidus necator*, *Dyadobacter fermentans*, *Pseudomonas stutzeri*, *Rhodobacter sphaeroides*, *Salinibacter ruber*, *Sulfurimonas denitrificans*, *Methylosinus trichosporium* Ob3b, *Methylocella tundrae*, *Methylomicrobium buryatense*, *Methanoregula boonei*, and *Methanolacinia petrolearia*. Strain information, including genome or fragment size and the number of target genes per organism are shown in Supplementary Table S1. DNA from cultured organisms was extracted from 1% SDS-treated cell pellets in CTAB buffer, followed by phenol:chloroform:isoamyl alcohol extraction and ethanol precipitation, as described before (Jones et al., 2013). The relative abundances of the microorganisms in the mock communities were multiplied with the median genome GC% contents to generate a weighted GC% content of the pool of samples with the following values: 47, 50, 53, 57, 60, and 63 (Supplementary Table S2). Pooling ratios of microorganisms’ DNA were calculated based on the expected number of functional genes in the extracted DNA, the DNA concentration (determined with a Qubit HS dsDNA kit (Thermo)) and genome size. Relative gene abundances in mock communities were calculated and compared to the sequences generated using targeted metagenomics. The relative abundance of each gene was calculated against the sum of abundance values of all 14 genes in the targeted metagenomes of the mock community samples (relative abundance of a specific gene = 100 x (gene reads / sum of reads of 14 genes)). The captured reads for mock community targeted metagenomics are shown in Table S4.

### Annotation of coding sequences from the mock community dataset

Predicted proteomes from the mock communities were retrieved from NCBI to count the number of each functional genes per genome. For *Nitrospira defluvii*; NC_014355.1, genes were predicted from the genome nucleotide sequence using Prodigal (v2.6.3) (Hyatt et al., 2010). Proteins were annotated using KOFAM (HMM database of KEGG Orthologs (KO)). Each protein was assigned the top hitting KO. Where available, cut-offs provided with KOFAMscan (Aramaki et al., 2020) were used to filter spurious results.

### Extraction of DNA and determination of chemical parameters from environmental samples

Samples from two different environments, an agricultural soil in Hungary (n = 3), and a wetland in Bellefontaine, France (n = 3) (Table S3), were collected in order to assess the effectiveness of the probe set in enriching functional genes in different sample contexts. Extraction of DNA from the environmental samples was performed as previously described by Siljanen et al., (2019). Briefly, samples were homogenized by bead-beating 0.5 g of soil at a speed of 5.5 m/s, for 30 sec with phenol:chloroform:isoamyl alcohol extraction in CTAB buffer, followed by ethanol precipitation. The quality of DNA extracts was assessed using a NanoDrop ND-1000 (Thermo), and DNA concentration was measured with a Qubit HS dsDNA kit (Thermo). The soil chemical parameters soil C/N ratio, pH, Fe II/III-, ammonium- and nitrate content were determined as described earlier (Siljanen et al., 2019; Bagnoud et al, 2020).

### Targeted metagenomic library preparation, target enrichment of libraries and sequencing with Illumina Miseq

To prepare DNA for targeted metagenomics with probe capture, DNA was first fragmented and indexed as follows. For each sample, sequencing indexes and sequencing adapters were provided as commercial service by the Centre for Genomic Research (CGR)-laboratories, at the University of Liverpool, Liverpool, UK. Libraries were produced with KAPA HyperPlus Library Preparation (Roche) kit to produce insert sizes of 630 bp according to the manufacturer’s instructions. The protocol is shown in Fig. 1. and described here briefly: 150ng of environmental DNA was fragmented with KAPA fragmentation buffer and KAPA Frag Enzyme at +37°C for 20min. Fragmented DNA was ligated to SeqCap Adapters A and B, +20°C for 15 minutes. Fragments were purified 0.65x volumes of AMPure XP Reagent for 5 min at room temperature in magnetic particle collector (MPC). Then the clear supernatant was removed and the library was washed twice with 200 μl of 80% ethanol. The libraries were removed from MPC by eluting them to 53 μl of elution buffer (10 mM Tris-HCl, pH 8.0) by incubated two minutes. Each library was amplified for 7 cycles, with KAPA HyperPlus Library Preparation kit in order to add the indexes for each sample (cycling conditions: +98°C, 15 sec; 60° for 30 sec; +72°C for 30 sec). Amplified libraries were purified with AMPure XP beads as above.

The hybridization reactions to the NimbleGen SeqCap EZ Developer Probe set were performed according to the manufacturer’s instructions. The hybridization was done in non-binding DNA background 25 μg, on top of 1 μg of each library equally pooled Multiplexed DNA Sample Library were added. The Multiplexed Hybridization Enhancing Oligo Pool (2,000 pmol in 2μl volume), the SeqCap HE Universal Oligo (1,000 pmol in 1μl volume) and (1,000 pmol in 1μl volume) SeqCap HE Index Oligo Pool) was mixed with 1 μg of equally pooled Multiplexed DNA Sample Library. Two volumes of AMPure XP Reagent was added on top of the above mixture and thoroughly mixed. Samples were incubated for 10 minutes to allow the sample library to bind to the beads. Samples were placed on the MPC to capture the beads and the solution allowed to clear. Once clear, the supernatant was discarded carefully, to not disturb the beads. Afterwards 190 μl of 80% ethanol was added to the samples containing the bead-bound DNA samples. The samples were left on the MPC during this step. Samples were incubated at room temperature for >30 seconds. The 80% ethanol was carefully removed and discarded, without disturbing the beads. Beads were dried at room temperature with the tube lid open for 5 minutes (or until dry).

A master mix of the following reagents was prepared, scaling up to reflect the number of captures: 7.5 μl of the 2x Hybridization Buffer, 3 μl of Hybridization Component A. Then 10.5 μL of the Hybridization Buffer/Hybridization Component A mix from the previous step was added to the bead-bound DNA sample. Samples were removed from the MPC and mixed thoroughly. It was important that enough mixing was performed at this step to yield a homogeneous mixture. It was left to sit at room temperature for 2 minutes. Sampled were then placed on the MPC. After liquid cleared, 10.5 μL of supernatant (entire volume) was removed and placed in a new tube containing 4.5 μl of the SeqCap EZ Developer Probe pool. This was mixed thoroughly. Hybridization incubation was performed in a thermocycler using the following program with heated lid set to 10°C above the block temperature: incubation was 95°C for 5min, and 47°C for 72 hours (SeqCap EZ HyperCap Workflow protocol, Manoharan et al., 2015).

To purify the hybridized libraries the SeqCap Pure Capture Bead Kit was used. To prepare the beads for binding the samples, 50 μl of beads (streptavidin coated magnetic beads) was added to the 1.5 ml tube at room temperature for capturing reaction. Tubes were placed on a MPC, the supernatant removed and beads were washed two times with 100 μl of 1x Bead Wash Buffer. Afterwards it was removed from the MPC and mixed by pipetting up and down. Beads were bound on the magnetic particle collector. Once the liquid was clear, the supernatant was removed. In the second wash, the beads were aliquoted with 50 μl of 1X Bead Washing Buffer. Beads were bound and the supernatant again removed. One hybridization sample (15 μl volume) was added to the washed SeqCap Pure Capture Beads. Samples were bound to the beads by placing the samples in a thermocycler at +47°C for 15 minutes (heated lid at +57°C). Then 100 μl of 1x Wash Buffer was added on top of the Capture Beads and bead-bound DNA and mixed thoroughly. The tube was placed on the MPC to capture the beads. The solution was allowed to clear, and the supernatant removed, while being careful not to disturb the beads. Afterwards, 200 μl of 1x Stringent Wash Buffer was added to each capture reaction. Those were mixed by pipetting up and down, when tubes were not in a MPC. Samples were placed in the thermocycler to +47°C for 5 minutes. The wash was repeated with 1x Stringent Wash Buffer. Next, 200 μl of 1x Wash Buffer I was added, vortexed for 10 second, and incubated 1 minute at room temperature. Once the solution was clear, it was removed, again being careful not to disturb the beads. The washing with 1X Wash buffer II was repeated, and then 1x with Washing Buffer III. Samples were removed from the MPC and 53 μl of PCR-grade water was added to the bead-bound sample. Sampled were then incubated at room temperature for 2 minutes and placed in the MPC. Then 50 μl of clear solution was collected in a new tube.

Sequencing for probe hybridized and washed DNA was performed with Illumina MiSeq PE300 chemistry in the Centre for Genomic Research (CGR), University of Liverpool, Liverpool, UK, resulting in 198,700-311,500 reads per sample for the environmental samples and up to 2,600,000 reads for the mock communities.

### Targeted metagenomics read processing and mapping

All six possible reading frames of the nucleotide sequence reads generated by the targeted metagenomic sequencing of the mock community samples were translated using transeq from the EMBOSS package (v6.6.0.0) (Rice et al., 2000). These were mapped to the genomes using DIAMOND blastp (v2.0.6.144) (Buchfink et al., 2015), with a minimum percentage identity of 60%. This threshold was selected as it resulted in the largest percentage of mapped reads while being stringent enough to prevent spurious mappings. No minimum coverage was used to account for reads that did not fully overlap with the proteins from the mock community dataset. For each direction, up to four matches were retained, to account for cases where a single read spanned neighboring proteins. Multiple matches in the same direction could overlap by up to 15 amino acids to account for possible protein fusions or sequencing errors. The blastx searches against the *refseq* database (Altschul et al., 1990) were performed in order to remove duplicates or reannotate falsely annotated entries due to the presence of homologous gene families in our dataset (*amoA* vs. *pmoA*, *nxrB* vs. *narH* nitrate reductase or *napA*+*narG* vs. *nrfA*). The blastx search was done to evaluate whether the read was correct or not, and the run was made with the default parameters of blast. Each of the (up to four) matches were used to assign KOs to the reads, based on the predicted KOs of the proteins to which they mapped. If, in a given direction, a read was assigned the same KO multiple times, this was collapsed into a single hit, retaining the hit with the best E-value.

### Evaluation of functional annotation of reads

Protein predictions for each read were performed using nhmmer from the HMMER suite (v.3.3)(Eddy 2011) using the in-house generated HMMs for target genes as well as KOFAM as described above. An inclusion threshold of <0.0001 was used when assigning protein predictions for the reads. For a given read, using all six of the translated frames, up to four protein assignments could be made, evaluated based on KO bit scores or identity filtering. Reads assigned KOs for the function of interest were counted as positively identified hits. Subsequently, reads were mapped to their corresponding coding sequences (CDS) from the mock community organisms to determine which reads constituted “true positives” (TP) or “true negatives” (TN). Reads mapped to a given CDS were assigned functions according to the above-mentioned KOFAM CDS annotation as their “true function”. If the HMM assignment (both in-house and from KOFAM) of a read matched the functional assignment of the CDS to which it was mapped, then it was counted as a TP. The TP, false positive (FP), and false negative (FN) values were summed up for all models using all methods, and precision and recall were calculated. Read mapping (described above) was used to determine the total number of possible TPs of the reads (Fig. S6), from which the actual number of TPs was subtracted to calculate the number of FNs. Precision was calculated as the ratio of TPs to total positive detections (i.e., number of TPs and FPs). Recall was calculated as the ratio of TPs to all theoretically obtainable true positives (i.e., sum of TPs and FNs). The script is available on Zenodo: https://doi.org/10.5281/zenodo.15752134;/probe-capture/tree/main/3-other_scripts/Angus_scripts-folder.

### Shotgun metagenomics library preparation, sequencing with Illumina HiSeq and analysis

Shotgun metagenomic sequencing was performed on environmental DNA (n = 3) from agricultural soil and wetland soil (Table S3), extracted as described above for TDB production. Illumina TruSeq library preparation with an insert size of 250bp was carried out with 1 μg of DNA and sequencing was performed in Illumina HiSeq PE150 lane as a commercial service in Vienna Biocenter Core Facility (VBCF, Austria) and Microsynth AG, Switzerland. In this approach, the previously generated HMM profiles were used to identify the target gene pools (E < 0.001). The predicted functions of each identified sequence were confirmed by tblastx against the *refseq* database, using DIAMOND (Buchfink et al., 2015). Annotation and identity information was further processed with awk in unix and in Rversion 3.5.3 to produce lists of community structures for each target gene. The relative abundance of each gene was normalized against the sum of total reads, Table S4: Relative abundance of gene x = (Abundance of gene x / total reads sequenced) x 100.

### *amoA* gene amplicon sequencing

Archaeal *amoA* gene amplicons from environmental DNA (n = 3) from the agricultural soil were generated as described earlier by Siljanen et al., (2019), and sequencing was performed with Illumina MiSeq PE250, in LGC, Münich, Germany. In brief, *amoA* gene amplicons were generated using 0.6 ng DNA template with GoTaq DNA polymerase (Promega) and 0.5 μM primers (CamoA_19F [5’-ATGGTCTGGYTWAGACG-3’] and TamoA_632R-4 [5’-GCKGCCATCCATCKRTANGTCCA-3’]) with a 5’-prime Illumina sequencing adaptor [5’-CTCTTTCCCTACACGACGCTCTTCCGATCT-3’] for both end of above primers for making the indexing possible in the sequencing service laboratory. Cycling conditions were the following: 1min initial denaturation at 95°C, 30sec denaturation at 94°C, 30sec annealing at 60°C and 45sec extension at 72°C, with 35 cycles. PCR reactions were performed in triplicate for each sample (n = 3), and purified with the High Pure PCR product purification kit (Roche). Sequencing indexes were added by LGC, where libraries were equally pooled and sequenced with Illumina MiSeq PE250.

### Processing of amplicon *amoA* reads

Archaeal *amoA* (Thaumarchaeal-*amoA*, *TamoA*) amplicon reads processing was carried out by a custom-based Python script available at Zenodo: https://doi.org/10.5281/zenodo.15752134;/probe-capture/tree/main/1-scripts/1-amplicon_seq_script_v1.sh-folder. Briefly, quality trimming of sequencing reads was performed by truncating amplicon forward read R1 reads to 200bp and reverse read R2 180bp, discarding all reads with an expected error greater than 0.5. The *amoA* reads were dereplicated and amplicon sequence variants (ASVs) were assigned to taxonomic bins described in the reference *amoA* gene database (Alves et al. 2018) using a 55% identity cutoff with USEARCH8 (Edgar, 2010). Chimeras were filtered out with UCHIME (Edgar, 2011). For the remaining ASVs, quality controlled R1 and R2 reads were fused together, classified with UCLUST in QIIME (Caparaso et al., 2010) and clustered to OTUs with 97% sequence identity cut-off for each taxonomic bins. Taxonomic bins of each *TamoA* gene cluster were generated using the reference gene database and taxonomy by Alves et al. (2018) (Altschul et al. 1990; Callahan et al., 2016). Relative abundance of *TamoA* taxonomic bins was calculated as follows: Relative abundance of *TamoA* taxonomic bin x = (Abundance of *TamoA* taxonomix bin x / total *TamoA* reads) x 100. In Fig. 4A, the relative abundance of each taxonomic bin (relative to the column sum) was plotted for each sample. The sequencing depth for the *TamoA* gene amplicon library was ∼27,000-58,000 PE250 reads per sample up to 31Mb for the agricultural soil (Table S4).

### Processing of metagenomic *amoA* reads

Archaeal *amoA* (Thaumarchaeal-*amoA*, *TamoA*) reads from targeted- and shotgun metagenomics, were identified with the *TamoA* in-house HMM model. The forward reads were dereplicated and assigned to the reference *amoA* gene database using a 55% identity cutoff with USEARCH8 (Edgar 2010). Chimeras were filtered out with UCHIME (Edgar 2011), all reads were clustered with UCLUST with 97% sequence identity in QIIME and assigned to taxonomic bins based on the closest *amoA* genes from the taxonomy database of OTUs (Alves et al. 2018) (Caporaso 2010). Relative abundance was calculated as follows: Relative abundance of *TamoA* taxonomic bin x = (Abundance of *TamoA* taxonomix bin x / total *TamoA* reads) x 100. In the targeted metagenomics approach, multiple detection of the same taxonomic bin is possible because of six probes are used for each sequence cluster. However, when relative abundance for each detected taxonomic bin is calculated for targeted metagenomic, if multiple detection occurs in the relative abundance calculation the community composition is balanced because of equal number of probes per each sequence cluster. Therefore, the relative abundance of different sequencing methods can be compared. The sequencing depth for the *TamoA* gene for three shotgun metagenomics *TamoA* reads out of ∼22-58M PE150 reads per sample up to 17 Gb, and for targeted metagenomics 4472-8230 *TamoA* reads out of ∼220-311k PE300 reads per sample up to 160 Mb, for the agricultural soil (Table S4). The metagenomic reads processing is described in the custom-made script available at Zenodo: https://doi.org/10.5281/zenodo.15752134; targeted with probe-capture: tree/main/1-scripts/2-probe_capture_script_v1.sh-folder, and shotgun metagenomics: tree/main/1-scripts/3-metagenomics_script_v1.sh-folder.

### Phylogenetic placement analysis of *nosZ* reads from shotgun and targeted metagenomics libraries

A reference alignment and phylogeny for *nosZ* was generated from full length *nosZ* amino acid sequences obtained from genomes downloaded from the NCBI genome database (accessed October 2019). A profile HMM for *nosZ* was generated from a previously published *nosZ* dataset, as described above (Graf et al, 2022). Reads were converted to amino acid sequences and then aligned using *hmmalign* within the HMMer suite (v.3.1.2)(Eddy 2011), and the homologous sections were identified using FastTree (v. 2.1.11, Price et al., 2010). The reference protein alignment was used for phylogenetic reconstruction using IQ-TREE (v1.6.12) with best model selection and 1000 ultrafast bootstraps. The Le-Gascuel (LG) substitution model with 10 rate categories was selected after automatic model selection. For the phylogenetic placement of *nosZ* reads obtained using either shotgun metagenome sequencing (n=3) or targeted metagenomics (n=3) of the agricultural site, the relevant reads were pooled together and clustered with CD-HIT (Fu et al., 2012) at 90% sequence identity before translating and aligning to the reference *nosZ* alignment using *hmmalign*. The translated reads were then placed in the reference phylogeny using the next generation evolutionary placement algorithm (EPA-NG v0.3.8; Barbera et al., 2019), with the same model parameters as used for constructing the reference phylogeny. The placement positions with the highest likelihood weight ratio were plotted on the reference phylogeny in R using the ‘ggtree’ package (v3.4.2, Yu et al., 2016). Script available for nosZ phylogenetic placements at Zenodo: https://doi.org/10.5281/zenodo.15752134; /probe-capture/tree/main/3-other_scripts/Chris_Jones_scripts-folder.

### Data and script availability

The shotgun metagenomic, targeted metagenomic and *amoA* amplicon sequencing data have been deposited in NCBI SRA under the Bioproject numbers PRJNA898102 and PRJNA488558. All scripts are available at Zenodo: https://doi.org/10.5281/zenodo.15752134. The repository contains as well the target gene database https://doi.org/10.5281/zenodo.15752134;probe-capture/2-selected_outputs/all_genes_compiled_ncbi_nt_envnt_fungene_split_file_0.fasta.gz-file (four files ..file_0-3.fasta.gz), and the final probe sequences https://doi.org/10.5281/zenodo.15752134;probe-capture/2-selected_outputs/Final_N_probes.list1.fasta-file.

## Results

### Benchmarking gene detection and quantification using targeted metagenomics in mock communities

The targeted metagenomic approach was evaluated using mock microbial communities comprising predefined mixtures of genomic DNA from 14 bacterial and three archaeal strains, together containing all 14 functional genes targeted by the probes and mixed in different proportions to produce six different median G+C contents ranging from 47% to 62% (Table S1). The overall relative abundances of target genes in the captured metagenomes across all GC% categories were correlated with their abundance in the original mock communities (*r* = 0.78 (*P* < 0.0001) (Fig. 2). Comparisons between the relative abundance of individual genes in the mock communities as compared to captured metagenomes revealed highly similar values (t = 1.8518e-10, df = 174.83, *P* > 0.05), with the exception of the *nirK* and the *hzoA* genes (Fig. S1a,b). Pearson correlation coefficients across all GC% categories varied between 0.76 and 0.93, although correlations were stronger within higher GC% categories (57%-63%; R = 0.80-0.93) than within lower GC% categories (47%-53%; R = 0.76-0.79) (Fig. S1). The probe capture targeted metagenomics produced 61-72% of target gene sequences from the total sequenced library of the mock-communities (Table S4).

**Fig. 2.**
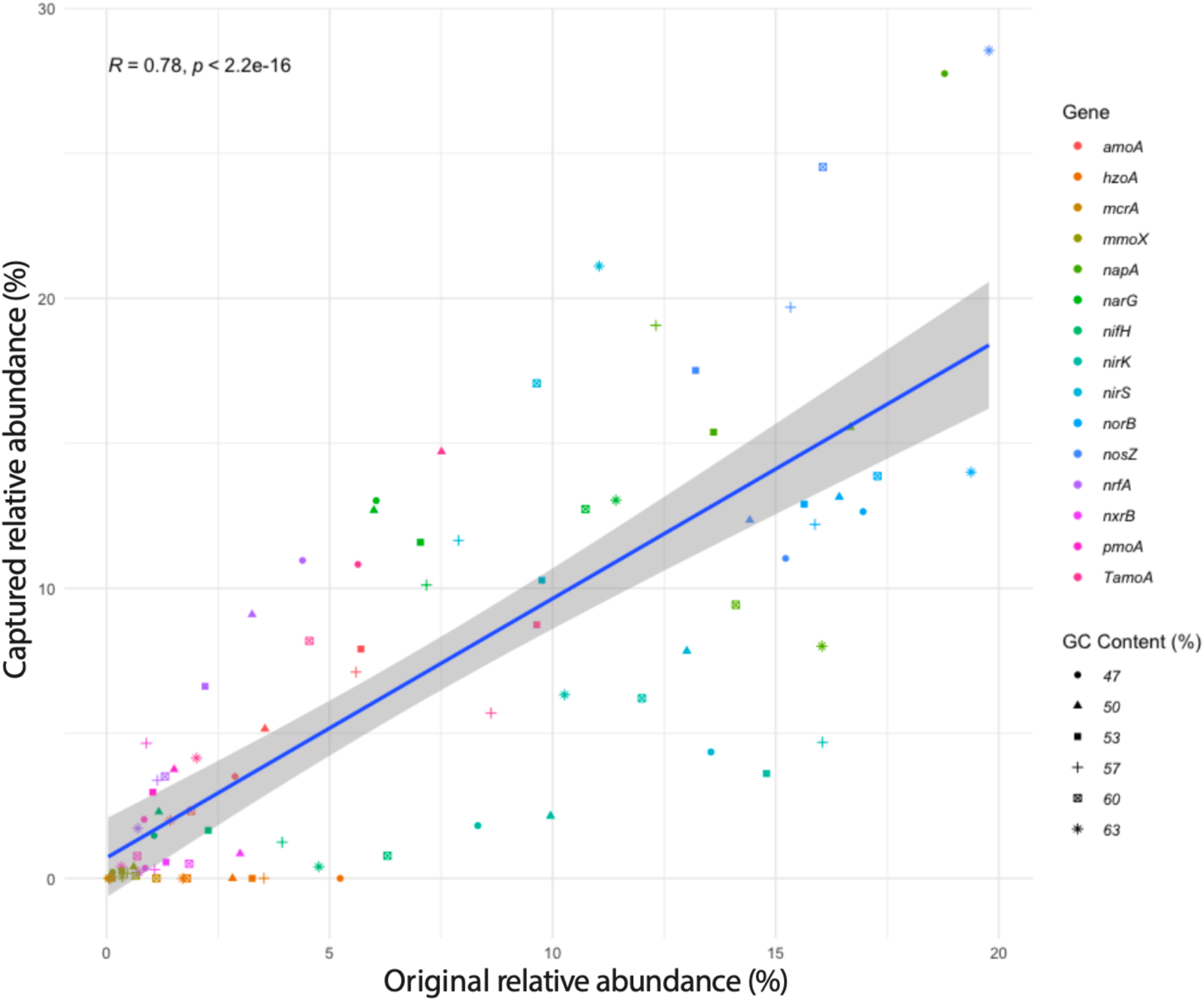
Comparison between the original composition (DNA of strains applied into the mock sample) of the mock community and the relative abundance for each functional gene detected using probe captured metagenomics for all GC% combinations combined. The relative abundance in the mock community was calculated based on the amount of each functional gene and organism as well as genome size. Color of the symbols depict gene and shape depicts the category of GC content (%).

### Precision of gene identification in mock communities

Precision for the in-house HMMs ranged from 74.1% (*nifH*) to 100% (*mcrA*) (Fig. S5). The average precision of the models was 93.4%, with a median of 99.8% (excluding *amoA* and *pmoA*). The recall ranged from 42.8% (*nxrB*) to 100% (*mcrA*). The average recall was 81.9%, with a median recall of 82.8%. Because KEGG only provides a single model covering the homologous genes *pmoA* and *amoA*, it was not possible to accurately calculate the precision and recall for these genes individually. A precision estimate could still be determined for the two genes, where a “true positive” is defined as a hit for a read identified by the in-house models as *amoA* or *pmoA* mapped to a CDS identified by the HMM for (*a*/*p*)*moA* genes provided by KEGG (K10944). In this case, the precision values for the *amoA* and *pmoA* genes were estimated to be 96.9%, and 100%, respectively. It is important to note that these estimates do not exclude the possibility that the in-house models misclassified reads belonging to these homologous protein families.

Precision and recall values were similar using KOFAMs instead of in-house HMMs. However, the results for *amoA* and *pmoA* genes were combined due to a single model for the associated KO for both genes, as mentioned above. Precision ranged from 76.5% (*amoA*/*pmoA*) to 100.0% (*mcrA*), while recall varied from 34.3% (*nifH*) to 100% (*mcrA* and *nxrB*). The average and median precision values were 94.5% and 97.7%, while the average and median recall values were 93.39% and 99.03%, respectively. The precision of KOs was lower for *napA* than for other genes when compared with custom DNA HMMs. This could be accounted for by the presence of formate dehydrogenase, a known homolog of *napA*. Formate dehydrogenase (K00123) was indeed identified among the genomes used for the mock community, so it is possible that the custom HMM was misclassifying reads originating from this known ortholog to *napA* (Cerqueira et al., 2015).

### Higher number of reads for all target functional genes but similar relative abundance obtained with targeted compared to shotgun metagenomics from complex environmental samples

We investigated the efficiency of the targeted metagenomics approach in natural complex ecosystems by directly comparing this approach with shotgun untargeted metagenomics generated with a nearly 100 times higher amount of sequencing (∼66 Mb and ∼6.4 Gb per sample, respectively) from wetland soils in Bellefontaine, France, and from an agricultural field in Hungary (Table S4). These soils represent distinct ecosystems with average physicochemical conditions, such as water content and oxygen concentration, and N availability that favors different distributions of functional genes (Jones et al. 2014). Despite the large difference in sequencing depth, targeted metagenomics generated a much larger set of target functional genes (Fig. 3). In both the agricultural and wetland site the targeted metagenomics approach detected 14 out of 15 gene-clusters and the shotgun approach 13 out of 15 genes (*mcrA* was not detected in the agricultural site and *hzoA* not in the wetland site). Moreover, up to 60 times as many identified gene sequences were detected using the probe capture compared to shotgun metagenomics (Fig. 3, Table S4). The relative abundances of functional genes in captured metagenomes correlated significantly with that in shotgun metagenomes from both the agricultural field (*R*_Pearson_ = 0.96, d.f.=14, *P* < 0.00001) and the wetland (*R*_Pearson_ = 0.70, d.f,=14, *P* < 0.01). The relative abundances of target genes were different between the two approaches (*P* > 0.00001) (Fig. 3). Moreover, increased relative abundance was detected with targeted compared to shotgun metagenomics in the agricultural site for *nifH*, *TamoA*, *nrfA*, *napA*, *nirK*, *nirS*, *norB*, *nosZ* and *mmoX* (Fig. 3a) genes and the wetland site for *nifH*, *napA, narG* and *nirS* (Fig. 3b) genes. The targeted approach generated a much higher number of reads for all target genes in both sites than shotgun metagenomics (Target reads in Table S4).

**Fig. 3.**
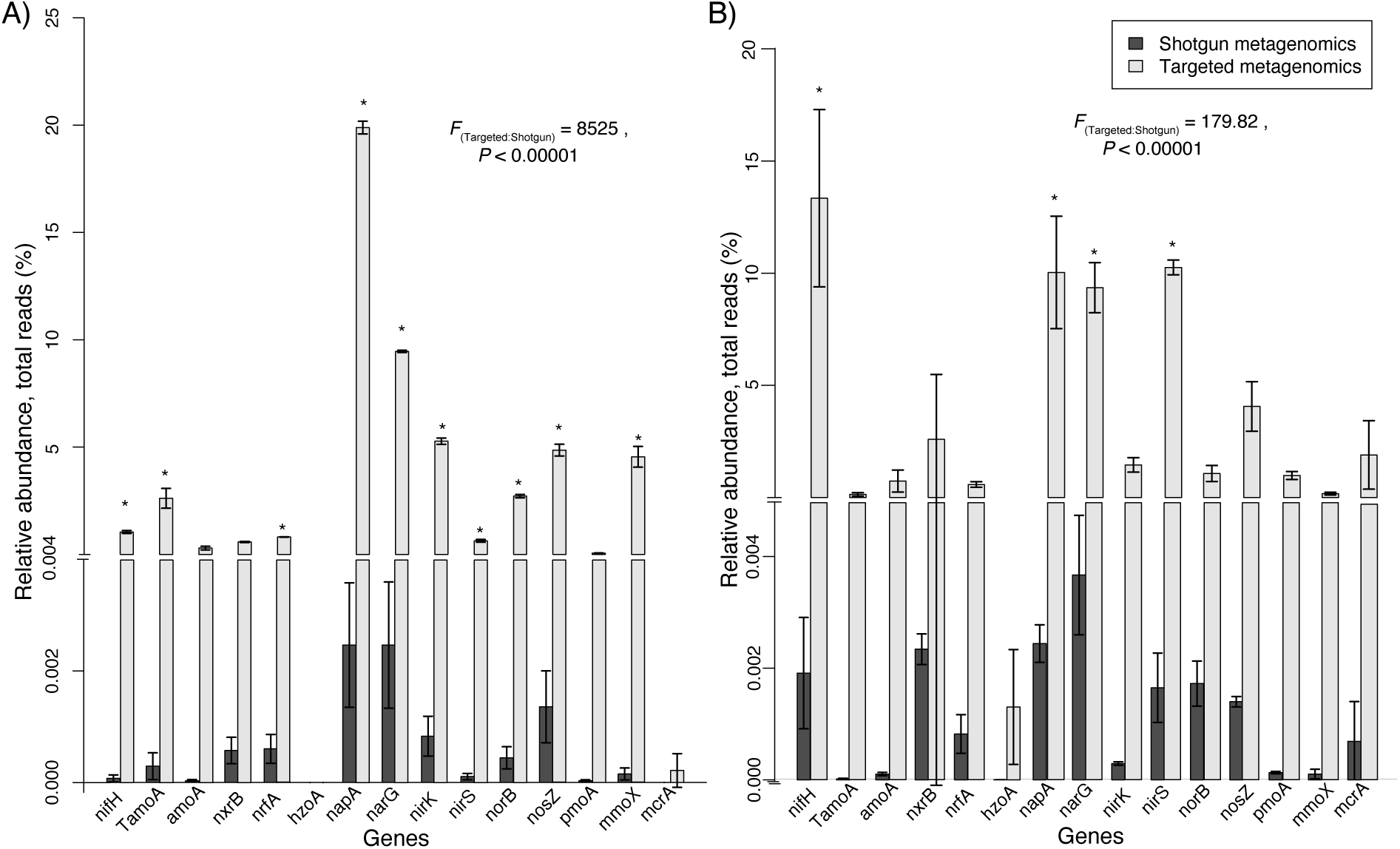
The relative abundance of functional genes (calculated against total sequencing reads) involved in inorganic nitrogen and methane cycling obtained by targeted metagenomics and shotgun metagenomics from A) agricultural soil; and B) wetland soil. Relative abundance was retrieved with HMMER searches of each functional gene. Statistically significant differences between shotgun and targeted relative abundances according to pairwise comparisons with ANOVA are shown with asterisk (*P* < 0.05, n = 3). The comparison of shotgun vs. targeted metagenomic measured with ANOVA is shown as *F* values and significance in the inset for both samples.

The different relative abundances of functional genes in the two ecosystems reflected known differences in their environmental conditions: (i) N_2_ fixation genes were expectedly higher in the unfertilized wetland; (ii) Genes involved in ammonia oxidation and nitric and nitrous oxide reduction were higher in the agricultural soil, which is likely to be more oxic and also subject to N fertilization which increase inorganic N cycling; (iii) Genes involved in methanogenesis were more abundant in the wetland, where anoxic conditions and methane production typically occur.

### Comparing diversity recovered for specific target genes from amplicon, targeted and shotgun metagenomics from complex environmental samples

In order to compare the diversity of specific functional genes detected by targeted metagenomics, shotgun metagenomics and PCR amplicon sequencing we focused on the archaeal *amoA* gene as a test case, as it is the second most sequenced gene in NCBI indicative of its ecological importance, and for which there is a comprehensive reference database for classification available (Alves et al., 2018). Moreover, archaeal ammonia oxidizers (AOA) typically outnumber their bacterial counterparts in most environments (Leininger et al. 2006), and even though they can be low abundant, their diversity is striking and underlines their distinct metabolic capabilities (Alves et al. 2018; Huang et al. 2021). Additionally, we compared the detection of *nosZ* genes between the targeted and shotgun metagenomics approaches in the agricultural soil, also based on an existing reference database (Graf et al., 2022). The following datasets were compared: (i) targeted and shotgun metagenomes, and PCR amplicons of the archaeal *amoA* gene in the agricultural soil (Fig. 4); (ii) *nosZ* gene sequences identified by targeted or shotgun metagenomics in the agricultural soil (Fig. S3).

Comparison of results from all three methods in agricultural soil samples showed that targeted metagenomics detected a substantially higher diversity of archaeal *amoA* genes than PCR amplicon sequencing, while, as expected, shotgun metagenomics detected a lower diversity than either of the other two methods (Fig. 4A). Specifically, targeted metagenomics detected 71-83 distinct taxonomic bins of *amoA* gene, whereas PCR amplicon sequencing and shotgun metagenomics detected 65-68 taxonomic bins and only three bins, respectively. Rarefaction analysis also indicated that targeted metagenomics retrieved a higher alpha diversity of archaeal *amoA* genes than PCR amplicon sequencing (Fig. 4B,C) and it also reaches a plateau much faster and with a lower number of sequences than amplicon sequencing (Fig. 4B).

Similarly, targeted metagenomes yielded a higher number of taxonomic bins for *nosZ* genes from the agricultural soil (∼6,800 bins) than shotgun metagenomics (∼1,300 bins) at a 90% identity cutoff, showing that the first approach detected a greater gene diversity with a much lower sequencing depth (Fig. S3). Gene sequence reads from both methods mapped to similar lineages in a comprehensive reference phylogeny of *nosZ* genes (Graf et al., 2022). However, a greater proportion of reads from the targeted metagenomics approach were assigned to deeper nodes of the phylogeny, particularly within the less studied clade II, suggesting that this method also captured more novel *nosZ* variants not represented in the reference phylogeny.

Targeted metagenomics (sequencing depth 117–160 Mb/sample) yielded 29.1±3.1 *TamoA* reads per 100,000 total reads, while shotgun metagenomics (7–17 Gb/sample) yielded only 0.0036±0.0039. The shotgun metagenomics produced only three quality-controlled *amoA* taxonomic bins were detected (Fig. 4A). Conversely, targeted metagenomics generated five times less *TamoA* reads (Fig. 4B) than PCR amplicon sequencing, with a sequencing depth of 15-31 Mb per sample. Nevertheless, the targeted metagenomics approach detected a much higher gene sequence diversity for both *TamoA* (Fig 4A, 4B) and *nosZ* (Fig. S3) than the two other methods in the agricultural soil samples. For *TamoA*, the targeted approach detected 28 times higher alpha-diversity than shotgun metagenomics, while 5 times higher diversity of *nosZ* genes was recovered, importantly covering genes affiliated with the less characterized Clade II. In turn, the targeted approach detected 1.24 times (24% more taxonomic bins) higher diversity of archaeal *amoA* genes than PCR amplicon sequencing in agricultural soils, respectively.

**Fig. 4.**
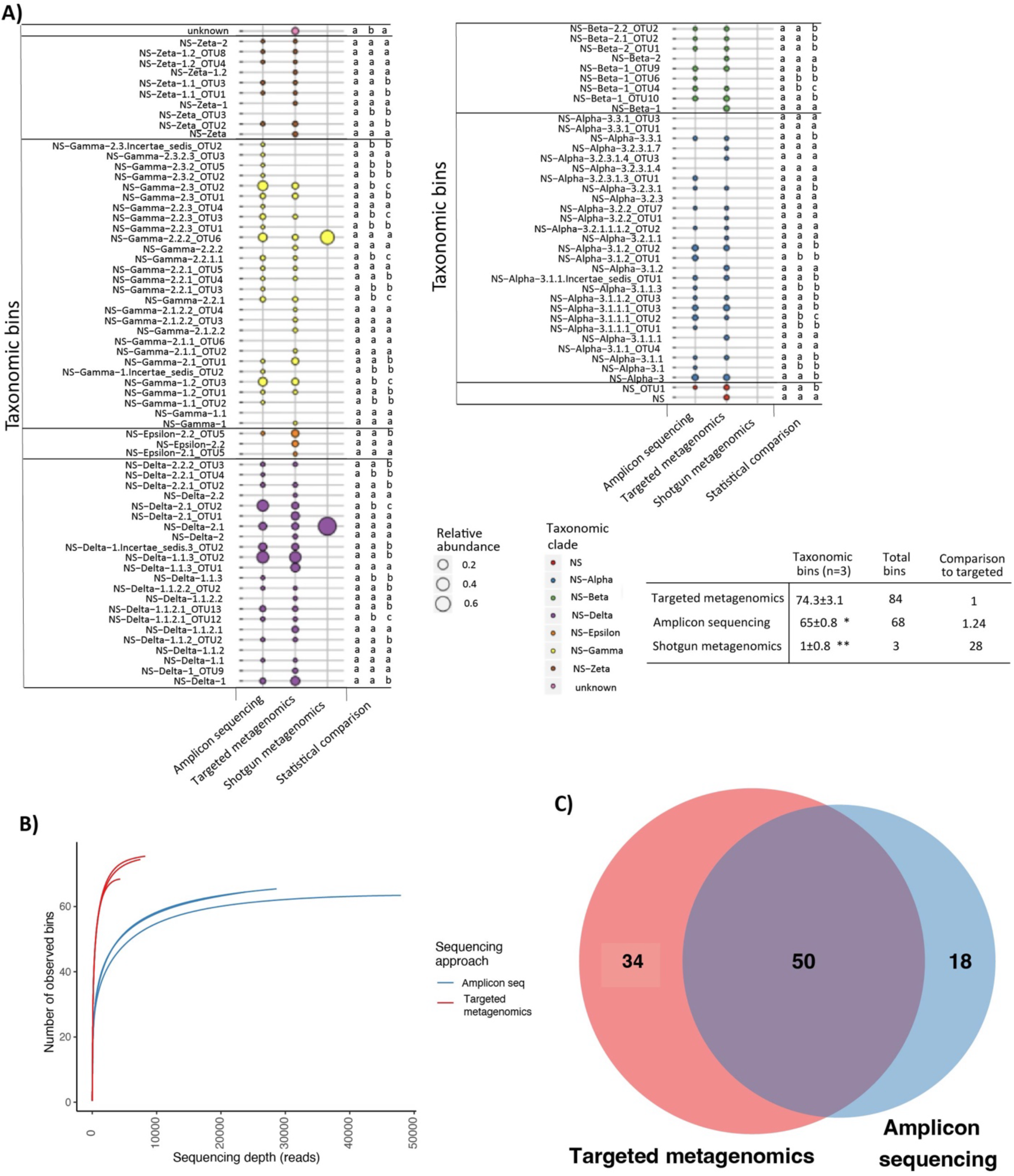
A) The relative abundance of archaeal *amoA* taxonomic bins produced from amplicon sequencing with *amoA* amplicon sequencing (*amoA* gene PCR amplicons sequenced with Illumina MiSeq, about 22000-47000 *amoA* reads per sample) (n = 3), metagenomics reads of targeted metagenomics (genomic DNA sequenced with Illumina MiSeq, after target enrichment, about 4500-8500 *amoA* reads out of 220000-310000 targeted reads per sample)(n = 3), or shotgun metagenomics (genomic DNA sequenced with Illumina HiSeq, about 22-57 million reads per sample, of which only two were archaeal *amoA*)(n = 3). In all cases, the analysis pipeline consisted of USEARCH comparison on *amoA* reference database (Alves et al., 2018) and taxonomy were analyzed with QIIME1 analytical pipeline. Sample having largest diversity is shown (n = 3). Bubble symbol size is proportional to the relative abundance of each taxonomic bin produced with sequencing approach in each sequencing method. Color indicates Nitrosospheria (NS)-clade of taxonomic bin. Statistically significant differences in abundance of each taxonomic bins between sequencing approaches based on Pairwise-Wilcox comparison are indicated with different letters (*P* < 0.05, n=3), and the significant difference of comparison for taxonomic bins is shown an asterisk, * *P*<0.05, ** *P*<0.01. B) The rarefaction analysis of archaeal *amoA* obtained by amplicon sequencing and captured metagenomics. C) Venn-diagram of archaeal *amoA* sequences obtained by amplicon sequencing and targeted metagenomics.

## Discussion

In this study, we report the development and evaluation of a new probe library for targeted metagenomics of 14 key genes involved in the microbial cycling of inorganic N and CH_4_. This included the compilation of databases containing all variants of the target genes from public databases, the development of an extensive set of probes targeting 14 well-established marker genes, and a bioinformatics pipeline to process and annotate the sequence data. This approach has been successfully used earlier to identify genes involved in microbial C metabolism in soil (Manoharan et al., 2015, Manoharan et al., 2017).

Our custom HMM models could assign reads to the correct genomes based on full protein sequences with higher precision than models based on KEGG Orthologs, especially when used to predict putative functions from short DNA fragments, such as metagenomic reads. This accuracy indicates that the HMMs can reliably classify reads in the mock community and are sensitive enough to detect distantly related genes or novel gene variants. Notably, reads identified as false positives may not actually be incorrect but could represent novel gene variants not captured by KEGG Orthologs.

In validation experiments with mock communities comprising 18 distinct organisms encoding multiple target genes, we showed that this method successfully reproduced the expected relative abundances of nearly all target genes, with the exception of the *nirK* and *hzoA* genes. Although 10.3% of the probes targeted *nirK* genes of nitrifiers, these were detected at a lower relative abundance than that present in the original mock community. However, the relative abundance of *nirK* genes matched the frequency of *nirK* genes only in heterotrophic organisms in the mock community, indicating a bias in the detection of distinct *nirK* gene variants. The sequencing technology used can influence the sequences generated due to known GC biases. In this case, the relative abundance of *nirK* genes may have been biased by the false low coverage of genomic regions with non-optimal GC% content (50-60%) known from MiSeq Illumina sequencing (Browne et al. 2020). Consistent with this observation, there was some variation in probe specificity among mock communities with different genomic GC% content, with high GC% communities having a generally higher target gene detection efficiency. For example, the mean GC% content of nitrifier *nirK* genes is lower than that of heterotrophic *nirK* genes (53% and >60%, respectively) and thus it is plausible that the later are sequenced more efficiently. This effect was also noticeable among *nosZ* genes, as the *nosZ* gene of *D. fermentas*, with GC content of 52.9% (Table S1, Fig. S1), was detected more efficiently than that of *S*. *denitrificans*, which has a 32.7% GC content. Moreover, variation in detection efficiency may also result from differences in probe hybridization. For instance, probes with higher GC% content are expected to bind more strongly to their targets given the higher binding energy between guanine and cytosine than between adenine and thymine. Although our approach successfully identified *hzoA* genes in the environmental samples, it failed to detect them in mock communities. It is possible that this was due to the PCR-generated genomic fragment used as source material for *hzoA* genes in the mock community, which, unlike other target genes provided in the whole genomic DNA, is more susceptible to degradation or structure-related constraints on probe hybridization.

Targeted metagenomics also closely reproduced the relative abundances of target genes obtained with shotgun metagenomics in two distinct natural microbial communities from soils. The relative gene abundances were strongly positively correlated between the two methods, indicating that the probe capture step did not introduce a significant quantitative bias. Importantly, this relationship held also true for *nirK* genes, confirming that the lower detection efficiency of nitrifier *nirK* genes was due to a variation in GC content. This observation emphasizes that particular attention should be given to the design of probes for low GC genes, such as by targeting gene regions that are not highly conserved among all gene variants, where higher GC probes are more likely to outcompete lower GC probes, as in the case of *nirK*. Nonetheless, as observed in the mock communities, the relative gene abundances quantified in environmental samples were strongly positively correlated between the two approaches to all GC% categories (Fig. S2). This further supports that the observed variation in detection efficiencies among gene variants with targeted metagenomics has a negligible effect on community profiles.

The thaumarchaeal *amoA* gene (*TamoA*) and the *nosZ* gene were used as case studies to investigate the differences in the detected diversity of single functional guilds that are known to be highly diverse (Alves et al. 2018, Graf et al. 2022). To determine the community composition of the *TamoA* gene for the different sequencing approaches, the reads are mapped to a reference library, in turn forming the taxonomic bins. Consequently, the same gene from the same organism can be detected multiple times in targeted metagenomics due to multiple probe hybridization. However, when calculation of relative abundance is performed, it balances the detected community because an equal number of probes is used per sequence cluster and therefore the relative abundance of targeted method can be compared to other sequencing tools. At first glance, PCR amplicon sequencing seems to have resulted in 18 more taxonomic bins than targeted metagenomics. However, these sequences were generated by a 35-cycle PCR, known to produce some chimeric sequences. Examples of such errors have been shown even for shorter amplification cycles (Kozich et al., 2013). Problems due to chimeric sequences in public databases from amplicon studies are known and were encountered during the compilation of the thaumarchaeal *amoA* reference database (Alves et al., 2018). Although we used standard pipelines for the detection of chimeric sequences, it is possible that we might have missed some, which may be represented in the 18 bins produced with amplicon sequencing, leading to an inflated number of sequences post-processing. The absence of long amplification steps in targeted metagenomics is expected to preclude the formation of chimeric sequences, which represents a significant advantage over typical PCR-based approaches.

It should be noted that the efficiency of the targeted metagenomic approach largely depends on the coverage and quality of the database used to generate the probes. We used hmmer profiles to collect all possible genes from the NCBI nucleotide and WGS databases, as well as Fungene and other published databases were used for the target gene library. Only one hmmer profile per gene was used for detection, however, clade specific hmmer profiles could also be used to have a broader outcome for neighbouring clades. We used fairly relaxed conditions when including thresholds for screening the NCBI databases, because we wanted to have closely related organisms and genes in the probe pool. For example, this was the case for the *mmoX* gene family, which also includes butane, propane and toluene monooxygenases. We included these closely related *mmoX* genes in the probes, therefore these closely related *mmoX* genes were found from coniferous trees (Putkinen et al., 2021). However, the database is not constrained to full-length gene sequences from a limited diversity of complete genomes and long metagenomic scaffolds available and the probe/target diversity can be greatly extended through the inclusion of gene fragments generated by PCR or from short metagenomic contigs. Despite its high target precision, this method may nevertheless capture non-target sequences, especially for orthologous gene families who have evolved different functions, as is the case for *napA* genes and formate dehydrogenases. Such cases emphasize the importance of thorough data annotation and filtering procedures. With the target gene database used to generate probes in this study, we managed to detect a higher diversity for the case study of *TamoA* genes than the shotgun sequencing approach, showcasing the advantage of “casting a wider net” with targeted metagenomics. Similar results were obtained in other targeted metagenomics applications, such as for detection of diverse resistome-virulome elements (Noyes et al. 2017) or for improved taxonomic microbial community characterization via 16S rRNA enrichment (Beaudry et al. 2021).

Despite the technological advances and decline in cost of high-throughput sequencing, shotgun metagenomics remain impractical and prohibitively expensive to capture the diversity of specific, low abundant functional groups in complex environments, such as soils and sediments, especially in longitudinal studies. The current cost of generating the probes for targeted metagenomics is about 20-50 € per sample depending on the probe manufacturer. If the goal is to have a focused, comprehensive view of the diversity of functional guilds involved in inorganic nitrogen or methane cycling in a certain ecosystem, then targeted metagenomics can circumvent the high cost and overabundance of data generated by shotgun metagenomics, as well as provide more quantitative data and more information on the diversity of the genes of interest. In turn, PCR-based assays are limited by the biases associated with any set of primers targeting a large diversity of sequences, assay efficiencies which preclude the detection of highly divergent new sequence diversity as well as known issues with the generation of chimeric sequences during amplification. The targeted metagenomics approach circumvents these limitations by using multiple short probes targeting different regions of each gene, which largely increases the likelihood of capturing sequence variants that elude PCR amplification. Thus, this approach has not only the potential to capture rare and novel gene diversity in complex environments, but also to identify cryptic microorganisms in low-biomass samples or involved in suggested CH_4_ metabolisms, such as in the tree phyllosphere (Putkinen et al., 2021) and nitrogen cycling in coral holobiont (Glaze et al., 2021). Moreover, targeted metagenomics can also overcome issues associated with running and comparing multiple independent PCR assays when investigating several distinct targets. In that sense, this approach effectively represents a PCR-independent, multiplex approach to characterize simultaneously and in-depth the distributions of a broad range of functional genes, providing a holistic view of the status of the nitrogen and methane cycles in the studied ecosystems. This is especially advantageous when combined with functional studies, such as the determination of N-transformation rates and *in situ* fluxes, as showcased by a study of N_2_O emissions in thawing Yedoma permafrost sites over time (Maruschak et al. 2021). In this study, the application of targeted metagenomics with the N-cycling probe dataset presented here revealed that changes in the N-cycling microbial community composition were responsible for an increase in N_2_O emissions in revegetated Yedoma soils, which had undergone thawing a decade prior.

Conclusively, the targeted metagenomics approach developed here provides an efficient and cost-effective strategy for studying microbial functional guilds that typically represent small fractions of natural microbiomes, and whose diversity is generally underestimated and highly underrepresented in metagenomic datasets. This approach also circumvents the limitations and biases associated with PCR-based methods and has higher potential to capture rare or novel functional gene diversity.

## Acknowledgements

H.S. was financially supported by the Atmosphere and Climate Competence Center (ACCC) Academy of Finland project number 337550, and Nitrobiome project 342362, in addition H.S. was supported by Saastamoinen foundation, FEMS Society and Niemi foundation. L.M., A.B., and C.S. were funded by ERC Advanced Grant TACKLE (695192), and R.J.E.A. was funded by project P25369 of the Austrian Science Fund (FWF). A.S.H and F.L.S were funded by ERC Starting Grant EvolPhisiol (grant agreement 803768). S.H and C.M.J were supported by the Swedish University of Agricultural Sciences (senior career grant 2019-2024). We thank Lea Wittorf for providing DNA extracts from isolates for the mock libraries.

## Authors contribution statement

H.S., C.S., R.A. designed study, H.S. collected the targeted gene library, R.A.,S.H.,C.J. provided reference libraries for *amoA* and *nosZ* genes, L.M. designed the probes, L.M., A.B., H.S. extracted the environmental DNA and did the shotgun metagenomics, H.S. and S.H. extracted and provided the mock sample cultured organisms, A.H., F.S. did the precision and recall analysis between the KEGG and HMM approaches, A.B., H.S. did the *amoA* diversity comparison, S.H.,C.J. did the *nosZ* diversity comparison. H.S. wrote the first version of manuscript and all co-authors further developed the text.

