## Supplemental material for "Targeted metagenomics using probe capture detect a larger diversity of nitrogen and methane cycling genes in complex microbial communities than traditional metagenomics"

a. National Bioinformatics Infrastructure Sweden (NBIS), SciLifeLab, Department of Laboratory Medicine, Lund University, Lund, Sweden.
b. Membratec S, Ecoprac de Daval C 1, CH-3960 Sierre, Switzerland.
   Climate and Ecosystem Sciences Division, Lawrence Berkeley National Laboratory, Berkeley, CA, USA

**Supplementary figures,**

A)

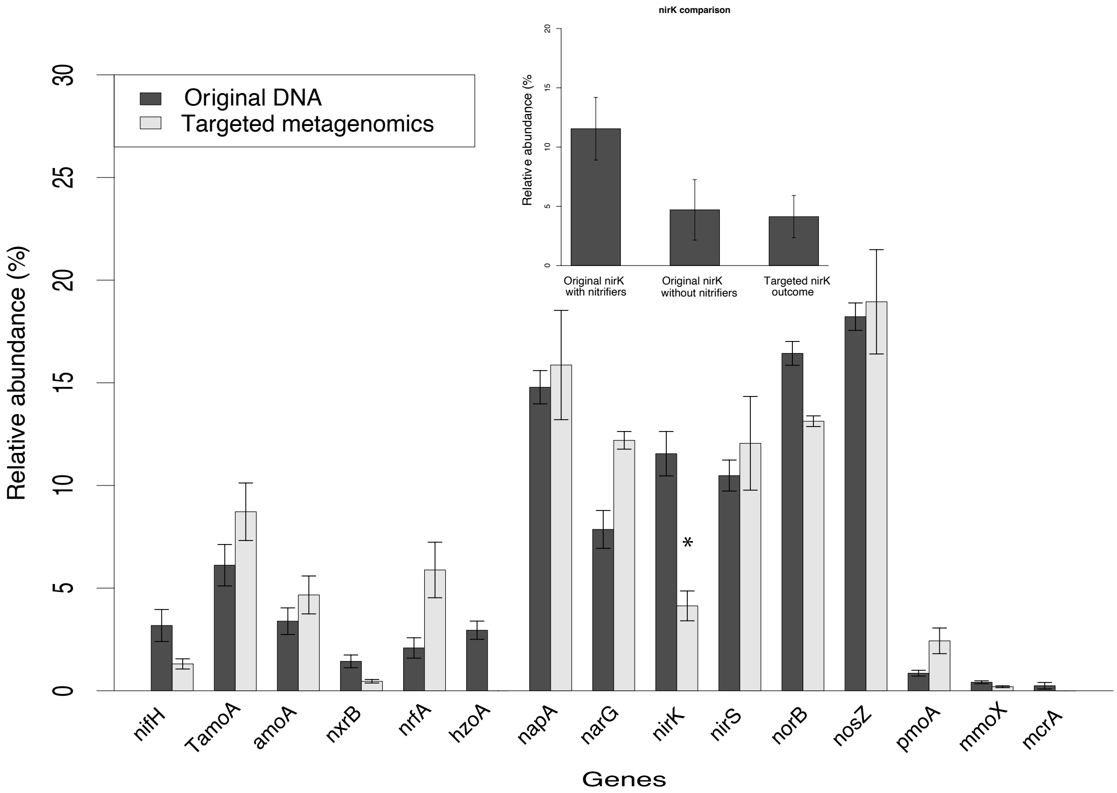


B)

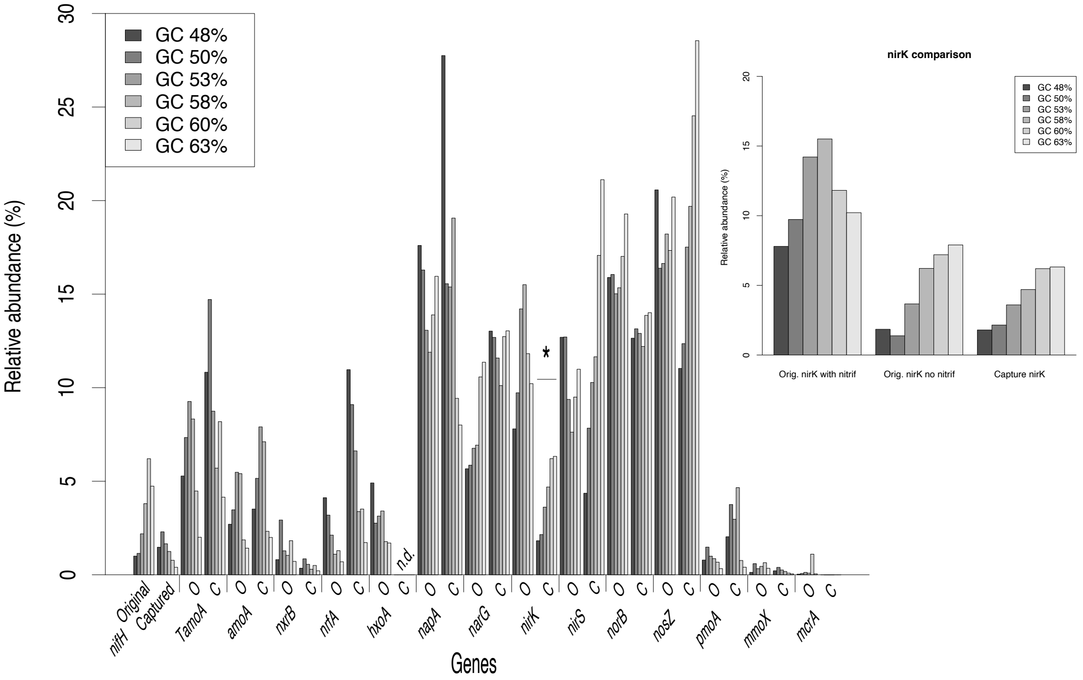


Fig. S1. The relative abundance of targeted functional genes originally mixed into the samples and in captured metagenomes retrieved with the custom HMMs searches. A) The relative abundance of each targeted functional gene in the original DNA mixture and that obtained by captured metagenomics (mean ± s.d., n=6) . Statistically significant differences between original and targeted metagenomic relative abundance according to pairwise comparisons with ANOVA are shown with asterisk ($P < 0.05$). B) Bars with different shades of grey show relative gene abundance values for each GC% category in the original community (O) and those obtained by targeted captured metagenomics (C).

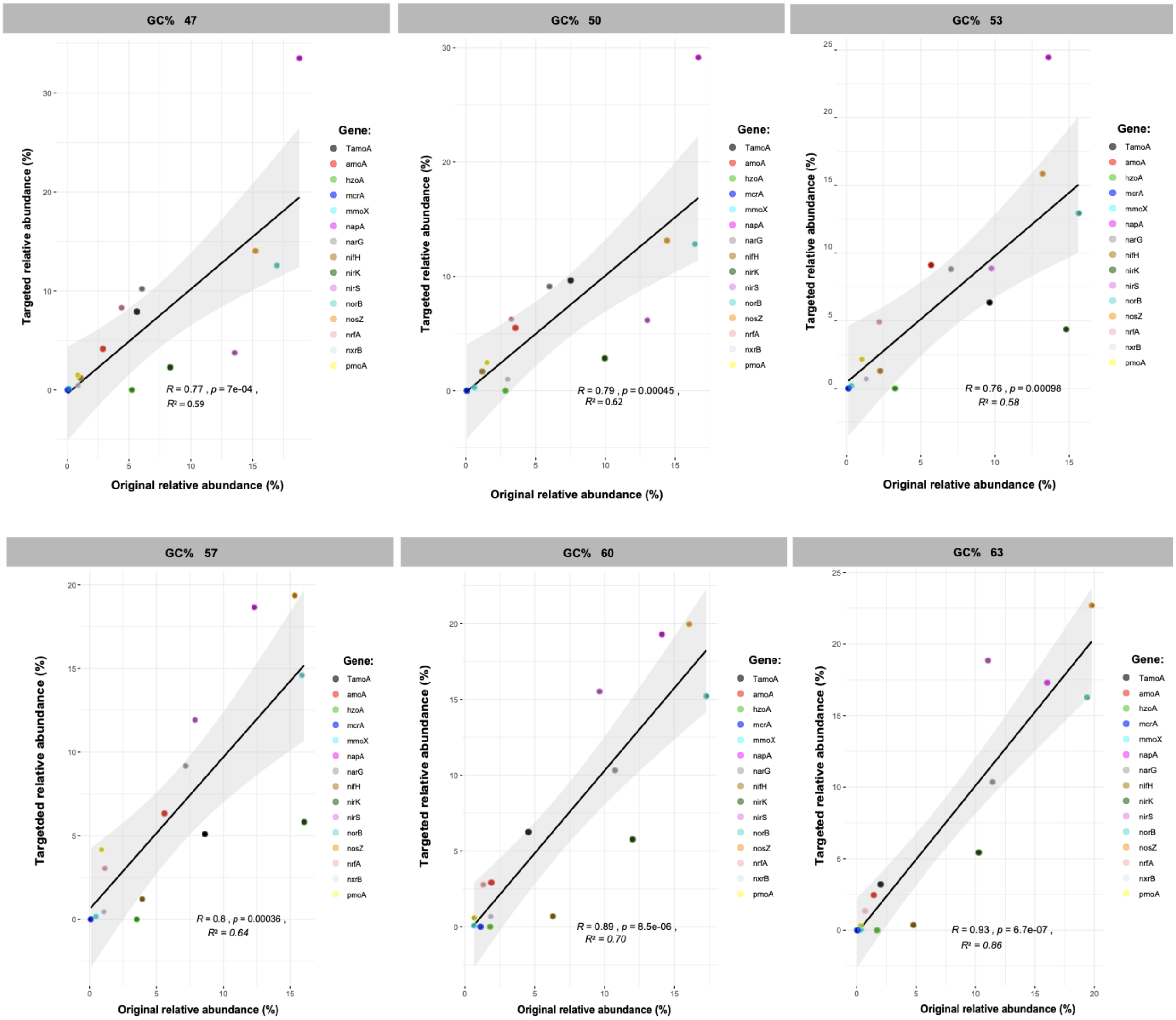


Fig. S2. Comparison of mock community between original (The relative abundance which was calculated with the amount of each functional gene and organism taken into the mock community and genome size.) and targeted metagenomics relative abundance produced for each different functional genes studied with probe hybridized targeted metagenomics for each GC% content separately.

Table S1. The mock-community organisms genome accession, presence of the N and $CH_4$ cycling genes in genome, median GC% of genome and DNA quality which was measured by for UV absorbance based on the with ratio of 260 [nm]/280 [nm] ratio.

| Organism: | genome accession (RefSeq) | Presence of N and $CH_4$ genes | Median GC% of genome | A260/A280 |
|---|---|---|---|---|
| *Nitrosospira multiformis* | NC_007614.1 | *amoA*, *nirK*, *norB* | 53.3 | 2.03 |
| *Nitrososphaera viennensis* | CP007536.1 | *TamoA*, *nirK* | 52.07 | 2.07 |
| *Nitrospira defluvii* | NC_014355 | *nxrB* | 59 | 1.27 |
| *Ca.* Kuenenia stuttgartiensis/*hzoA PCR fragment from plasmid vector* | | *hzoA* | 46.02 | 1.88 |
| *Pseudomonas aeruginosa PA96* | CP007224.1 | *narG*, *napA*, *nirS*, *norB*, *nosZ*-I | 66.2 | 2.11 |
| *Escherichia coli DH5a* | AE014075.1 | *nrfA*, *narG*, *napA* | 50.6 | 1.92 |
| *Shigella sonnei* strain FC1706 | CP014099.2 | *nrfA*, *napA* | 50.7 | 2.05 |
| *Cupriavidus metallidurans* CH34/CCUG 13724 | NC_007973.1 | *nosZ*-I, *nirS*, *norB*, *narG*, *napA* | 63.58 | 2.15 |
| *Cupriavidus necator* ATCC 17699 | NC_008313.1; NC_008314.1 | *nosZ*-I, *nirS*, *nor*, *narG*, *napA* | 66.3 | 2.16 |
| *Dyadobacter fermentans* DSM 18053 | NC_013037.1 | *nosZ*-II | 51.5 | 2.05 |
| *Pseudomonas stutzeri* JM300/DSM 10701 | NC_018177 | *napA*, *narG*, *nirS*, *nirK*, *norB*, *nosZ*-I | 63.3 | 2.17 |
| *Rhodobacter sphaeroides* DSM 158/ATCC 17023 | NC_007493.2; NC_007494.2 | *nifH*, *napA*, *nirK*, *norB*, *nosZ*-I | 68.77 | 2.16 |
| *Salinibacter ruber* DSM 13855 | NC_007677.1; NC_007678.1 | *nosZ*-II | 65.98 | 1.9 |
| *Sulfurimonas denitrificans* DSM 1251 | NC_007575.1 | *nosZ*-II, *nirS*, *norB*, *napA* | 34.5 | 2.13 |
| *Methylosinus trichosporium Ob3p* | CP023737.1 | *nifH*, 3x *pmoA*, *mmoX* | 65.82 | 1.42 |
| *Methylocella tundraea* | GCF_900749825.1 | *nifH*, *mmoX* | 63 | 1.11 |
| *Methylomicrobium buryatense 5B* | FO082060.1 | 1x*pmoA*, *norB* | 48.07 | 2.00 |
| *Methanoregula boonei* | NC_009712 | *nifH*, *mcrA* | 54.5 | 1.43 |
| *Methanolacinia petrolearia* | NC_014507 | *nifH*, *mcrA* | 47.4 | |

Table S2. The mock-community relative abundance composition, and weighted GC% content of each microorganisms and on each GC% category samples.

| **Organism:** | **GC 47% [Rel. Abun.]** | **GC 47% Weighted GC%** | **GC 50% [Rel. Abun.]** | **GC 50% Weighted GC%** | **GC 53% [Rel. Abun.]** | **GC 53% Weighted GC%** | **GC 57% [Rel. Abun.]** | **GC 57% Weighted GC%** | **GC 60% [Rel. Abun.]** | **GC 60% Weighted GC%** | **GC 63% [Rel. Abun.]** | **GC 63% Weighted GC%** |
|---|---|---|---|---|---|---|---|---|---|---|---|---|
| *Nitrosospira multiformis* | 0.024 | 1.30 | 0.034 | 1.82 | 0.053 | 2.80 | 0.05 | 2.79 | 0.02 | 1.18 | 0.02 | 0.98 |
| *Nitrososphaera viennensis* | 0.143 | 7.43 | 0.22 | 11.27 | 0.267 | 13.88 | 0.24 | 12.60 | 0.16 | 8.31 | 0.08 | 4.04 |
| *Nitrospira defluvii* | 0.011 | 0.65 | 0.043 | 2.55 | 0.018 | 1.09 | 0.02 | 0.89 | 0.03 | 1.92 | 0.01 | 0.82 |
| *Ca.* Kuenenia stuttgartiensis/hzoA fragment in plasmid vector | 0.13 | 6.10 | 0.0 | 3.75 | 0.090 | 4.16 | 0.10 | 4.56 | 0.06 | 2.91 | 0.07 | 3.02 |
| *Pseudomonas aeruginosa* PA96 | 0.004 | 0.28 | 0.008 | 0.52 | 0.014 | 0.93 | 0.013 | 0.83 | 0.026 | 1.72 | 0.03 | 1.83 |
| *Escherichia coli* , DH5α | 0.096 | 4.84 | 0.084 | 4.28 | 0.048 | 2.44 | 0.023 | 1.14 | 0.041 | 2.05 | 0.02 | 1.15 |
| *Shigella sonnei* strain FC1706 | 0.016 | 0.79 | 0.010 | 0.49 | 0.013 | 0.65 | 0.009 | 0.48 | 0.005 | 0.28 | 0.004 | 0.21 |
| *Cupriavidus metallidurans* CH34/CCUG 13724 | 0.012 | 0.78 | 0.027 | 1.72 | 0.045 | 2.84 | 0.048 | 3.04 | 0.091 | 5.80 | 0.11 | 6.70 |
| *Cupriavidus necator* ATCC 17699 | 0.019 | 1.23 | 0.029 | 1.90 | 0.043 | 2.85 | 0.054 | 3.60 | 0.090 | 5.99 | 0.15 | 9.91 |
| *Dyadobacter fermentans* DSM 18053 | 0.17 | 8.78 | 0.068 | 3.50 | 0.114 | 5.86 | 0.099 | 5.08 | 0.054 | 2.80 | 0.02 | 1.02 |
| *Pseudomonas stutzeri* JM300/DSM 10701 | 0.022 | 1.42 | 0.025 | 1.58 | 0.045 | 2.82 | 0.064 | 4.05 | 0.129 | 8.16 | 0.13 | 8.46 |

| | | | | | | | | | | | | |
|---|---|---|---|---|---|---|---|---|---|---|---|---|
| *Rhodobacter sphaeroides* DSM 158/ATCC 17023 | 0.021 | 1.46 | 0.011 | 0.79 | 0.045 | 3.12 | 0.092 | 6.34 | 0.111 | 7.61 | 0.17 | 11.35 |
| *Salinibacter ruber* DSM 13855 | 0.021 | 1.39 | 0.029 | 1.928 | 0.050 | 3.294 | 0.117 | 7.704 | 0.114 | 7.554 | 0.171 | 11.26 |
| *Sulfurimonas denitrificans* DSM 1251 | 0.26 | 9.85 | 0.287 | 9.889 | 0.123 | 4.260 | 0.043 | 1.475 | 0.002 | 0.073 | 0.008 | 0.28 |
| *Methylosinus trichosporium* Ob3p | 0.001 | 0.07 | 0.005 | 0.343 | 0.003 | 0.188 | 0.003 | 0.206 | 0.006 | 0.387 | 0.003 | 0.21 |
| *Methylocella tundraea* | 0.002 | 0.16 | 0.012 | 0.777 | 0.007 | 0.412 | 0.010 | 0.627 | 0.017 | 1.078 | 0.010 | 0.65 |
| *Methylomicrobium buryatense* 5B | 0.018 | 0.88 | 0.028 | 1.346 | 0.020 | 0.967 | 0.016 | 0.752 | 0.006 | 0.308 | 0.003 | 0.16 |
| *Methanoregula boonei* | 0.001 | 0.028 | 0.001 | 0.059 | 0.002 | 0.103 | 0.001 | 0.050 | 0.019 | 1.039 | 0.001 | 0.06 |
| *Methanolacinia petrolearia* | 0.000 | 0.011 | 0.001 | 0.024 | 0.001 | 0.042 | 0.001 | 0.040 | 0.010 | 0.479 | 0.000 | 0.02 |
| sum: | 1.00 | 47.43 | 1.00 | 48.51 | 1.00 | 52.70 | 1.00 | 56.26 | 1.00 | 59.64 | 1.00 | 62.10 |

Table S3. Soil properties of used samples in the shotgun vs. targeted analysis and targeted vs. archaeal *amoA* amplicon analysis.

| Site | Coordinates | Corg/Norg | pH $H_2O$ | $NH_4^+$ [µg $NH_4^+$ -N·g -1 dry soil] | $NO_2^-$ [µg $NO_2^-$ -N·g -1 dry soil] | $NO_3^-$ [µg $NO_3^-$ -N·g -1 dry soil] | Fe II [µmol·g -1 dry soil] | Fe III [µmol·g -1 dry soil] | Ref.: |
|---|---|---|---|---|---|---|---|---|---|
| Agricultural field, Hungary[#&%] | 47°31N 16°59E | 17.9±5.3 | 6.7±0.16 | 0.02±0.004 | 0.23±0.1 | 3.4±1.9 | 1.5±0.07 | 43.3±4.8 | This study |
| Belfontaine wetland, France[#] | 46°34N 6°04E | 14±0.6 | 6.9±0.1 | 0.04±0 | 0.04±0 | 0.002±0.001 | 78.5 ± 43.9 | 78.5 ± 43.9 | Bagnoud et al., 2020 |

bd = below detection limit

NA = not analyzed

#= shotgun vs. targeted comparison, Fig. 2,3.

&= Targeted vs. *amoA* amplicon analysis, Fig. 4, S3.)

Table S4. Number of total sequences produced, targeted reads and recovery of targeted sequences after capture and depth of the sequencing.

| Sample: | Sequencing method | Total reads | Targeted reads | % Targeted from total | Depth, of bases: |
|---|---|---|---|---|---|
| Mock community GC 47% | Targeted metagenomics | 1844716 | 1128755 | 61.2 | 1.0G |
| Mock community GC 50% | Targeted metagenomics | 2281740 | 1437919 | 63.0 | 1.3G |
| Mock community GC 53% | Targeted metagenomics | 2006493 | 1360701 | 67.8 | 1.1G |
| Mock community GC 57% | Targeted metagenomics | 2346193 | 1692037 | 72.1 | 1.3G |
| Mock community GC 60% | Targeted metagenomics | 2625812 | 1771389 | 67.5 | 1.5G |
| Mock community GC 63% | Targeted metagenomics | 2429736 | 1640046 | 67.5 | 1.3G |
| 44 FR_Bellfonte_wetland_BF_3_8 | Targeted metagenomics | 214947 | 124886 | 58.1 | 114.4M |
| 50 FR_Bellfonte_wetland_BF_10-1 | Targeted metagenomics | 285234 | 161399 | 56.6 | 147.5M |
| 51 FR_Bellfonte_wetland_BF_10-2 | Targeted metagenomics | 198756 | 109938 | 55.3 | 104.1M |
| 47 HG_agricultural_soil_H16 | Targeted metagenomics | 246209 | 130449 | 53.0 | 132M |
| 48 HG_agricultural_soil_H17 | Targeted metagenomics | 220005 | 116976 | 53.2 | 117.5M |
| 49 HG_agricultural_soil_H18 | Targeted metagenomics | 311535 | 162396 | 52.1 | 160.7M |
| Bellfonte_wetland BF_3-8 | Shotgun metagenomics | 21464245 | 5102 | 0.024 | 6.4G |
| Bellfonte_wetland BF_10-1 | Shotgun metagenomics | 43620482 | 6434 | 0.015 | 12.9G |
| Bellfonte_wetland BF_10-2 | Shotgun metagenomics | 50203331 | 6865 | 0.014 | 14.9G |
| agricultural_soil H16 | Shotgun metagenomics | 53422156 | 3457 | 0.007 | 16G |
| agricultural_soil H17 | Shotgun metagenomics | 22140941 | 2883 | 0.013 | 6.6G |
| agricultural_soil H18 | Shotgun metagenomics | 57814232 | 4213 | 0.007 | 17.3G |
| Agricultural H16 | Amplicon sequencing | 58006 | 49679 | 85.64 | 31M |
| Agricultural H17 | Amplicon sequencing | 27565 | 23648 | 85.79 | 14.9M |
| Agricultural H18 | Amplicon sequencing | 35955 | 30830 | 85.75 | 19.3M |

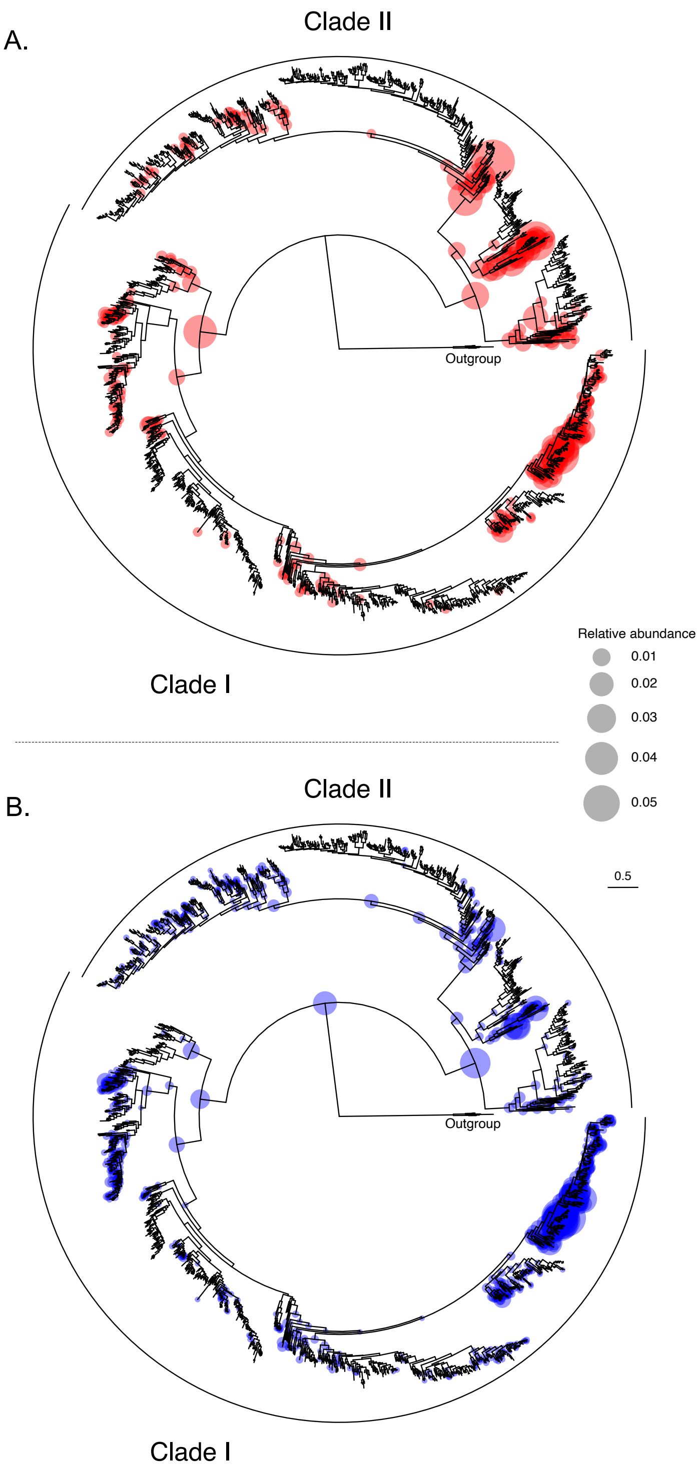


Fig. S3. Phylogenetic placement of *nosZ* reads obtained from agricultural soil using (n=3) A) shotgun metagenomics and B) targeted metagenomics (n=3). For illustration purposes all the replicas are pooled together. The reference phylogeny (Graf et al., 2022) is based on amino acid sequences analyzed using the LG+R10 substitution model in IQ-TREE, and node symbols indicate the location of placements in the reference tree. Symbol size corresponds to the relative abundance of reads placed at each node, and the scale bar indicates branch length in the reference tree. ‘Outgroup’ denotes distant homologues of *nosZ* with unknown function.

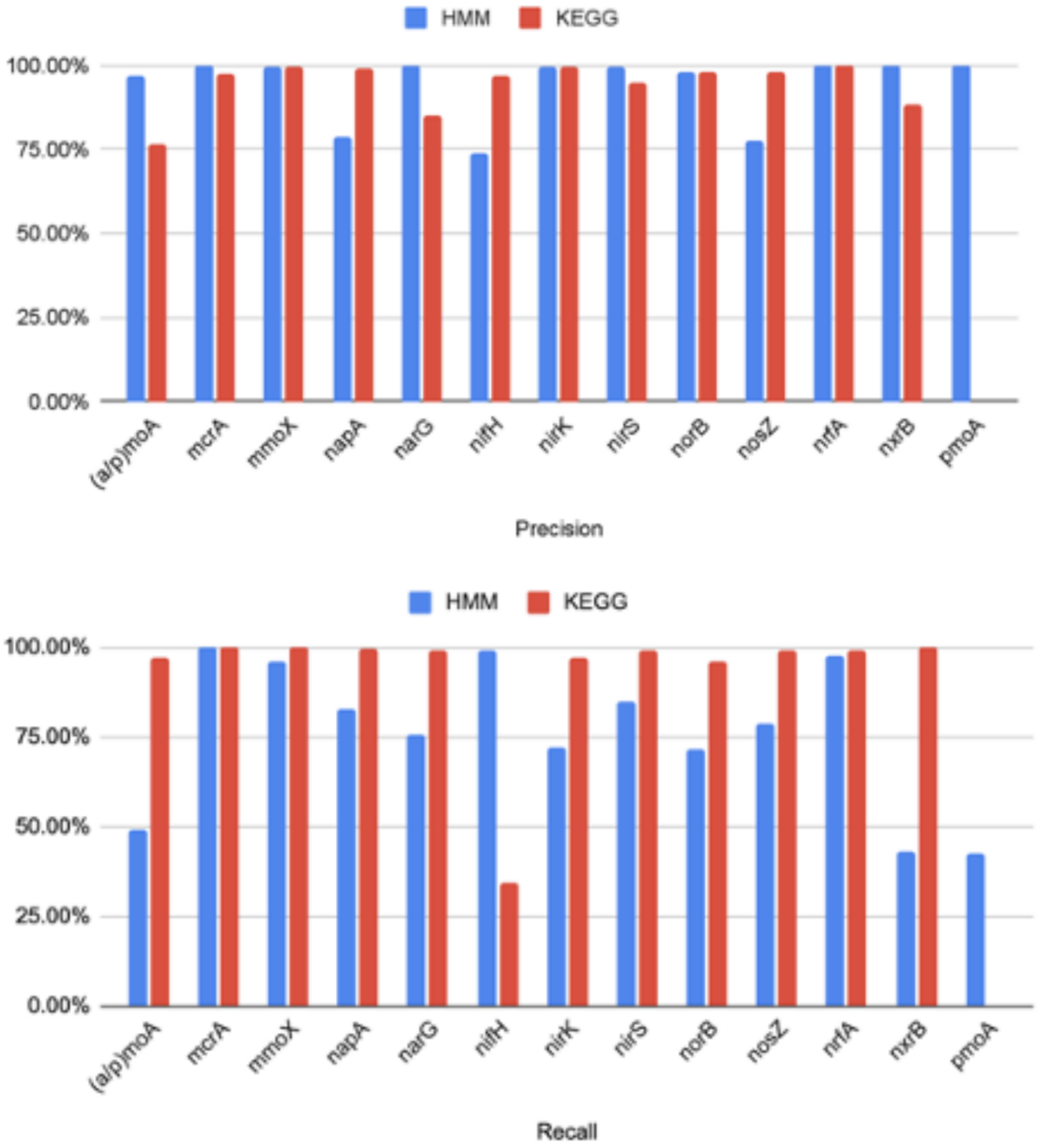


Fig. S4. Comparison of KEGG annotation and custom HMMs precision and recall for the mock-community comparison with different GC% categories combined.

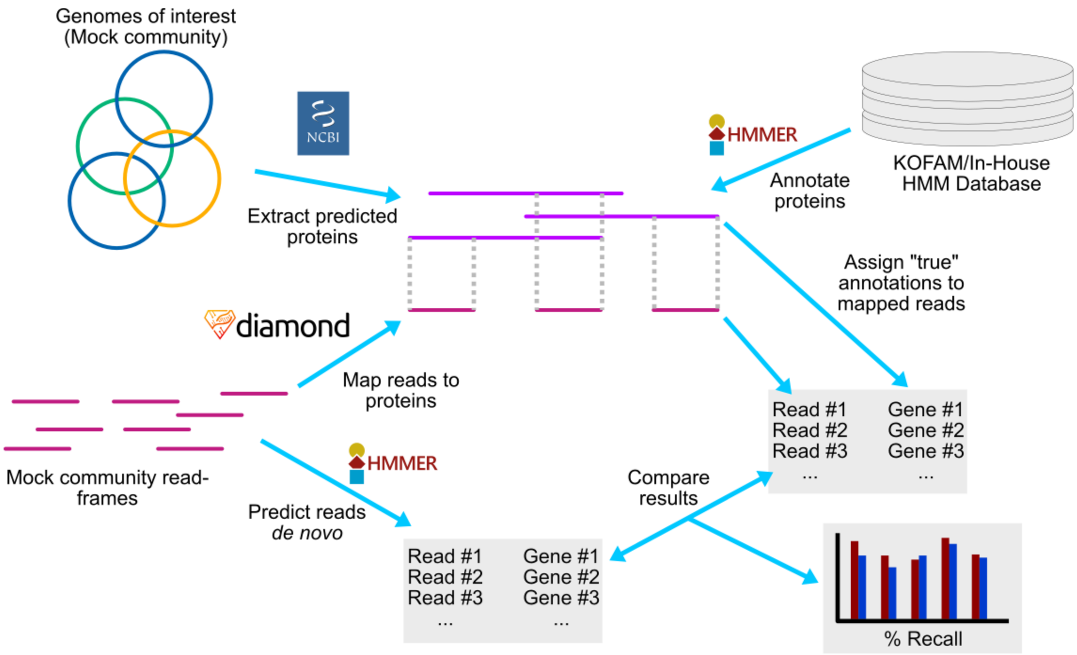


Fig. S5. Illustration of pipeline for determining TP, FP, TN, and FN de novo predictions by read mapping and HMM model prediction compared to genome predicted proteins.
